# Optical-quality assessment of a miniaturized intraocular telescope

**DOI:** 10.1101/2022.12.02.518911

**Authors:** Irene Nepita, Raffaele Raimondi, Simonluca Piazza, Alberto Diaspro, Faustino Vidal-Aroca, Salvatore Surdo, Mario R. Romano

## Abstract

**Purpose:** Evaluating the optical transmission and geometrical aberrations of an intraocular device, namely, the Small-Incision New Generation Implantable Miniature Telescope (SING IMT™, Samsara Vision), designed to correct age-related macular degeneration.

**Methods:** Optical transmission in the spectral range 350-750 nm of the implantable optics was recorded with a fiber-optic spectrometer. Geometrical aberrations were studied by measuring the wavefront of a laser beam after passing through the implantable optics and performing an expansion of the measured wavefront into a Zernike polynomial basis. The study was conducted under *in-vitro* experimental conditions. A second monofocal intraocular lens (SY60WF, Alcon) was tested and used as reference for assessing the optical quality of the SING IMT™ device.

**Results:** Spectroscopy measurements revealed that the SING IMT™ and monofocal IOL element feature UV-rejection and blue-rejection capabilities, respectively. Wavefront concavity indicated that the SING IMT™ behaves as a diverging lens with a focal length of approximately -100 mm; Zernike analysis showed that SING IMT™ has negligible coma, trefoil, astigmatism, and spherical aberrations of any order and along any direction.

**Conclusions:** The SING IMT™ exhibited even optical transmission in the whole visible spectrum and curvature capable of magnifying the retinal images without introducing geometrical aberrations, which proves the feasibility of this device as high-quality optical element for imaging. The rigidity of the compound lens of the SING IMT™ prevents mechanically-induced distortions, an issue encountered with polymeric lenses.

**Translational Relevance:** Spectrometry and *in vitro* wavefront analysis provide evidence supporting the new generation miniaturized telescopic intraocular lens as a favorable option to intraocular implant in age-related macular degeneration.

## Introduction

Age-related macular degeneration (AMD) is the leading cause of blindness in millions of people worldwide.^1,2^ Briefly, AMD strikes the center of the retina, namely the macula, leading to blurry and poorly resolved vision. So far, there is no effective treatment for advanced stage AMD. As a result, advanced stage AMD patients are forced to use supportive measures for hypo vision such as external magnifiers or telescopes. Surgical options, on the other hand, are very limited and include macular translocation surgery and more recently, retinal-pigmented epithelium transplant.^3^ In this scenario, the development of an effective intraocular implantable telescope for AMD may fulfill a considerable unmet need.

Recently, a miniaturized intraocular optical device consisting of a compound lens equipped with foldable haptics has been proposed (Samsara Vision Inc.), which can be surgically implanted into the eye through a preloaded delivery system.^4-7^ Once inside the eye, the implant creates a magnified image of the viewed object on the healthy portion of the retina. The healthy portion of the retina surrounds the degenerated macula because only the center of the retina is damaged in AMD disorders. The implant enables the retinal cells surrounding the macula to image the object.

The aim of the present study is the *in-vitro* optical quality assessment of the SING IMT™ device. To this end, we measured the geometrical aberrations and the spectral transmittance of the SING IMT™ optical system. Furthermore, we used a second mono-focal implantable lens (Monofocal IOL, +20 Diopters, Model SY60WF, Alcon), widely-used in cataract surgery, as a benchmark for the optical quality of the SING IMT™ device. We found that the SING IMT™ has an excellent optical transmission in the visible with negligible geometrical aberrations, which make this intraocular device a valuable tool for vision-improving applications.

## Methods

### Transmission spectra measurement setup

Measurements of the transmission spectra of the intraocular lenses were carried out by means of an optical fiber setup (Figure 1A)^8-9^. In particular, a wideband (350-750 nm) radiation of a tungsten halogen lamp (6V/30W, OSRAM) was shed against the element under test through a multimode optical fiber. The lenses under test were mounted into specific sample holders (Figure S1-2) and inserted into the optical path. The light passing through the optics under test was launched into a multimode optical fiber by means of a converging lens (LA1213, Thorlabs) and recorded with a spectral resolution of 0.38 nm through a photospectrometer (USB200+, OceanOptics). To increase the signal-to-noise ratio of our measurements, the recorded spectrum was averaged over 10 acquisitions, each integrated over a time interval of 50 ms.

**Figure 1:**
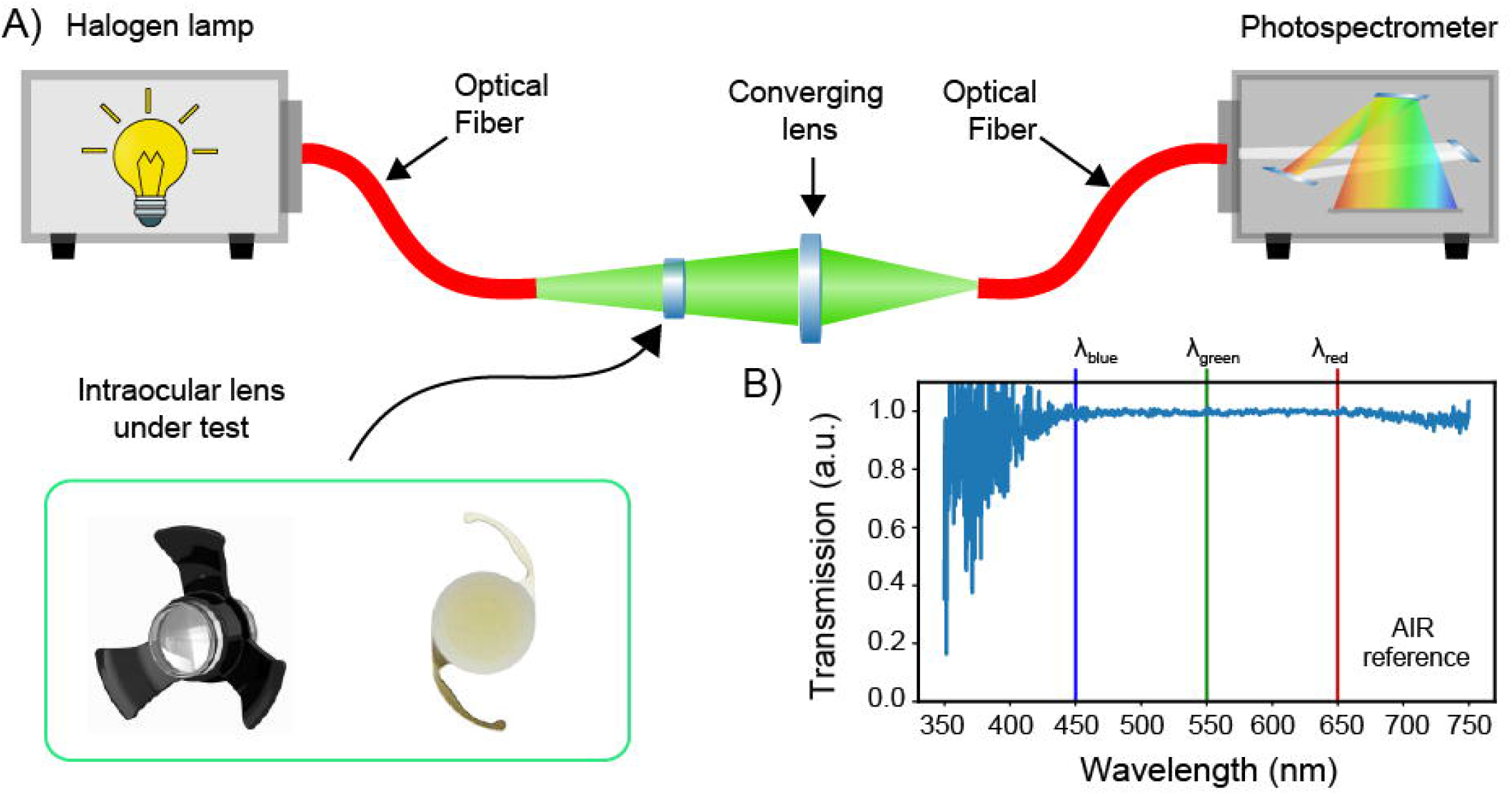
Optical transmission spectroscopy. A) Schematic description of the optical fiber setup used for measuring the transmission spectra of the intraocular optics. B) Reference spectrum of an ideal transmitter measured in air without any intraocular optical elements in the optical path.

A calibration procedure was performed to compensate for the spectral signature of the lamp and to remove possible background noise. To this end, we first subtracted the dark spectrum, namely the transmission spectrum with the lamp off, from the recorded spectrum and then we normalized the latter to the spectrum of an ideal transmitter, namely air (Figure 1B). As such, all the subsequent transmission spectra were measured as variations caused by the element under test with respect to air. Finally, each spectrum was normalized to its maximum, thus neglecting optical losses due to the limited coupling efficiency between the transmitted light and the collecting optical fiber.

The following analytical parameters were extracted from the measured transmission spectra: i) the transmission bandwidth defined as the full width at half-maximum (FWHM) of the transmission spectra; ii) the band flatness computed as the standard deviation of the transmission values within the FWHM; iii) the extinction ratios of the blue radiation, by definition the ratio between the transmission at the wavelength of 450 nm and the transmissions at 550 and 650 nm, respectively.

### Geometrical aberrations measurement setup

Geometrical aberrations of the intraocular optical devices were calculated by means of wavefront sensing and Zernike’s expansion, an approach widely used in ophthalmology and optical metrology.^10-11^ In particular, we designed and assembled a custom setup based on free-propagation of a laser beam and comprising two main functionalities: i) preconditioning of the laser beam, and ii) wavefront sensing with a Shack-Hartmann sensor.

The beam conditioning setup (Figure 2A-C) was designed and arranged in order to obtain a laser beam with the following properties: Gaussian shape, collimation, and a beam waist filling the optical aperture of the intraocular device. To this end, a continuous-wave (CW) laser emitting at 530 nm (Coherent, Sapphire) was filtered and expanded with an afocal 4f-system and a 50-μm pinhole (Figure 2A). The collimation of the laser beam was verified with a shear plate (SI050, Thorlabs) (Figure 2B). A fraction of the beam was redirected into a CMOS camera (DCC1545M, ThorLabs) to compute its size through beam profiling methods (Figure 2C). The beam waist at 4-sigma was 3.2 mm was in agreement with the clear aperture of the SING IMT™ device. For the measurements with the monofocal IOL, which has an optical aperture of 6.5 mm, an additional afocal system with a 2×-magnification was used to further expand the laser beam. The remaining fraction of the beam was directed towards a Shack-Hartmann sensor (WFS150-5C, Thorlabs) to compute the wavefront distortions and the Zernike polynomials (Figure 2D).

**Figure 2:**
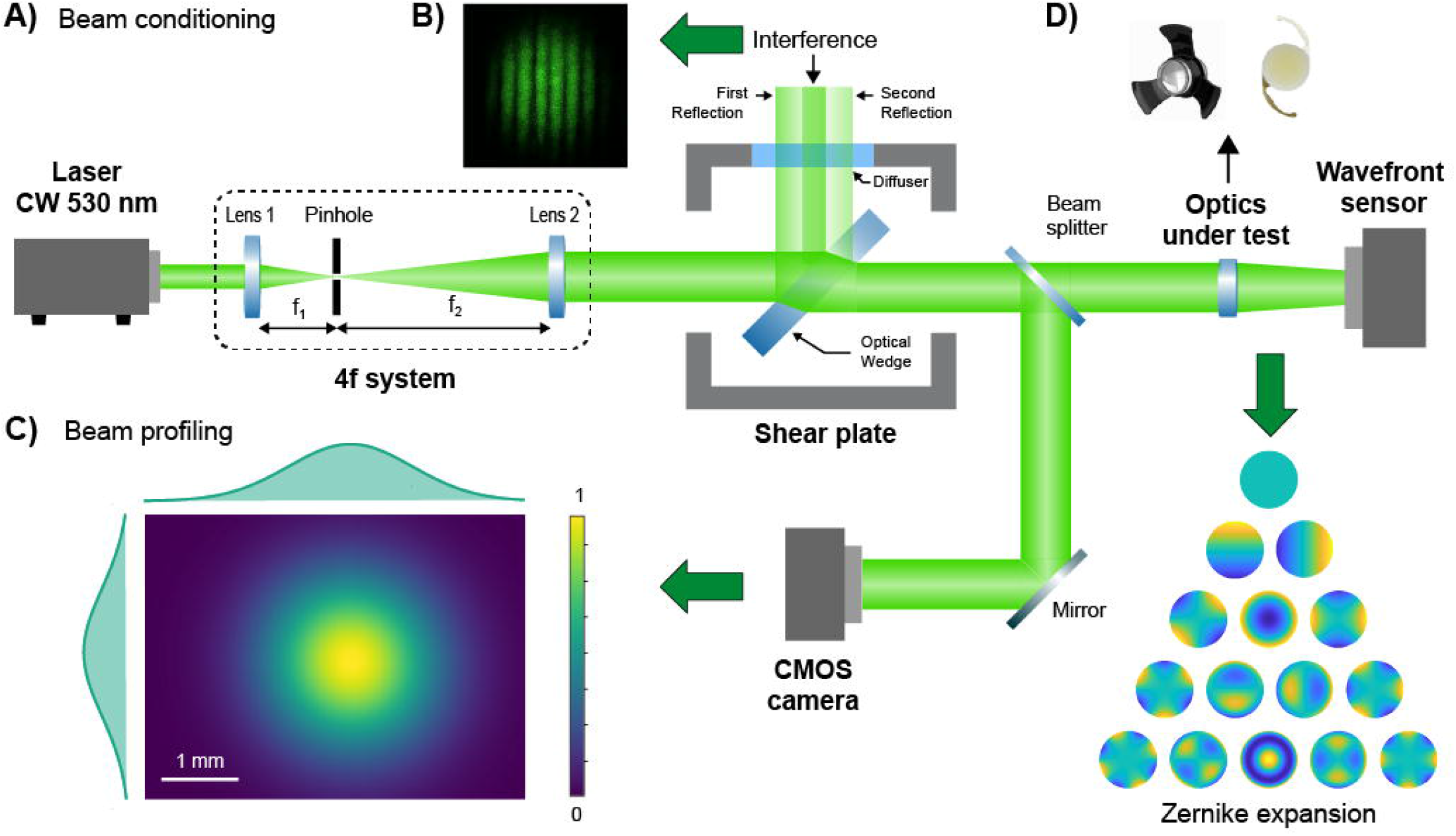
Wavefront sensing setup. A) The beam exiting a laser module is filtered and expanded with a 4f-system and a pinhole. B) A shear plate is used to set the distance between the lenses of the 4f-system thus ensuring collimation. C) A fraction of the laser beam is imaged and processed with a beam profiling algorithm in order to determine the beam waist. D) The reaming fraction of the beam is directed towards the intraocular lens under test and the resulting wavefront measured and expanded in Zernike polynomials.

In order to only measure the wavefront distortion due to the optical element under test, a calibration was necessary. Precisely, we recorded the wavefront of the preconditioned laser beam only and used it as a reference (Figure S2A). The reference wavefront was substantially flat (Figure S2B) with no evident optical aberrations (Figure S2C). Once calibrated, the optical setup allowed us to precisely measure the wavefront distortions due to any optical elements inserted in the light path. To ensure the reliability of our measurements, each wavefront was computed by averaging 10 acquisitions and 3 wavefronts recorded for each lens. Moreover, in order to provide high resolution to our measurements, namely accurate detection of a large number of Zernike terms, we selected a pupil diameter (i.e., 3.3 mm) close to the beam waist. For experiments with the SY60WF monofocal IOL, we fulfilled this requirement by inserting a 4f-system with a magnification of 0.5× between the intraocular lens and wavefront sensor. The contribution of these additional optics to the wavefront was subtracted during calibration.

## Results

### Transmission properties

Figure 3 shows the normalized transmission spectra of the SING IMT™ and SY60WF monofocal IOL. We found that the IOL device largely attenuates the intensity of light in the blue band of the visible spectrum (Figure 3A). The SING IMT™ device, on the other hand, evenly transmits the entire visible light, slightly attenuating the blue band below the 400 nm and entirely stopping the UV components (Figure 3B).

**Figure 3:**
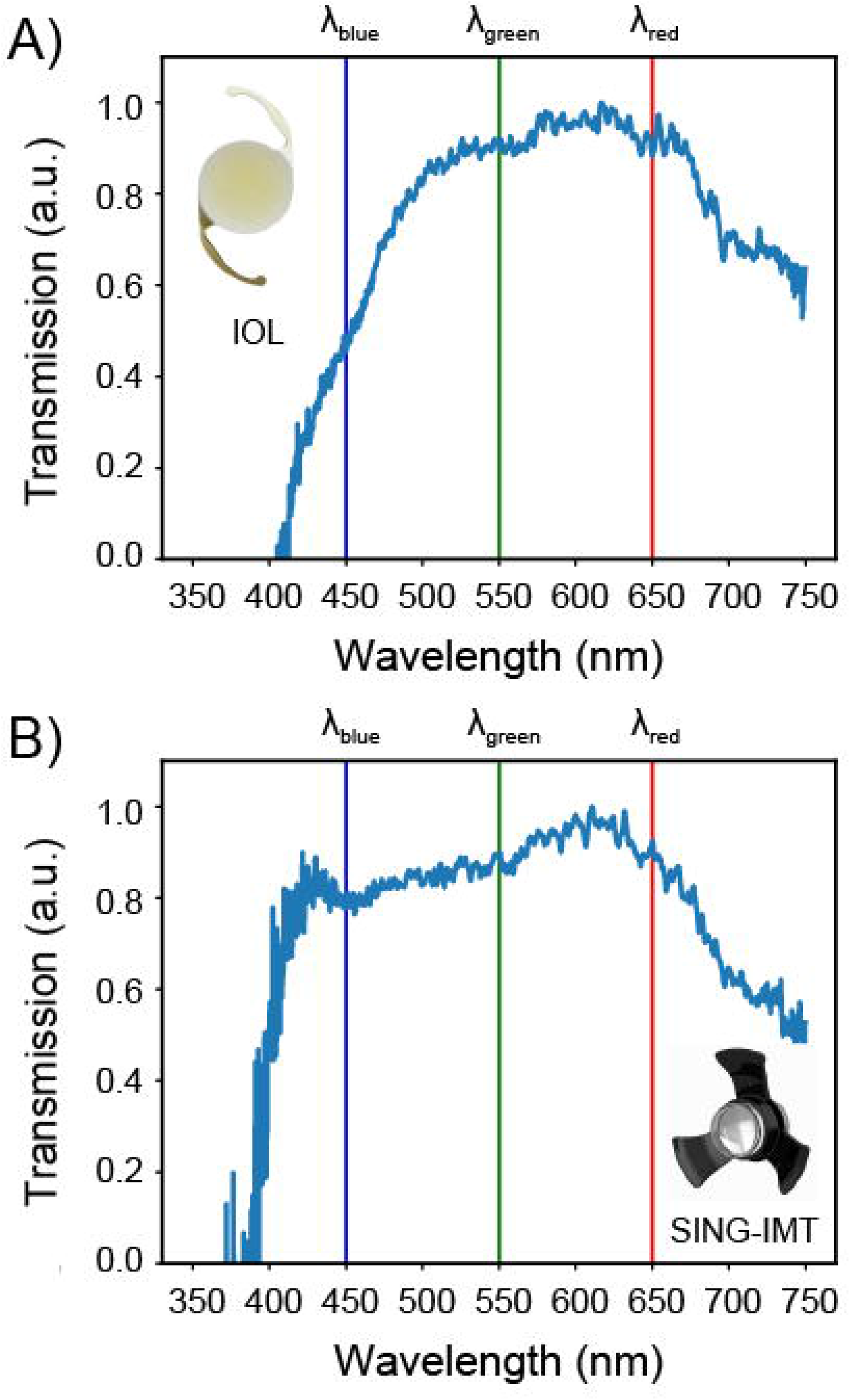
Transmission properties in the UV-Vis. Experimental normalized transmission spectrum of the SY60WF monofocal IOL IOL (A) and SING IMT™ device (B). The vertical blue, green, and red lines mark the wavelength values of 450, 550, 650 nm.

In order to quantify these effects, we extracted several analytical parameters from the recorded spectra, precisely both width and flatness of the transmission band along with the extinction ratio between transmitted light intensity at the wavelengths of 450, 550, and 650 nm. Results of this analysis are presented in Table 1. The same parameters are reported for the case of the ideal transmitter, namely, air. We found that both devices substantially exhibit a flat response within their bandwidth. However, the bandwidth of the SY60WF monofocal IOL is 56 nm narrower than that of the SING IMT™. The calculated extinction ratios confirmed the different spectral behavior of the two devices. Indeed, the SY60WF monofocal IOL shows an attenuation of the blue light, with respect to the green and red, of approximately 50%. The SING IMT™, on the other hand, has almost near-unity extinction ratios and no filtering capabilities in the blue spectrum, as per optical design.

**Table 1:** Main features of the recorded transmission spectra for an ideal transmitter (i.e., air), the SING IMT™ optics, and the SY60WF monofocal IOL, respectively.

|  | Bandwidth (inf-sup)<br>(nm) | Flatness<br>(rms) | Extinction ratio<br>(blue to green) | Extinction ratio<br>(blue to red) |
| --- | --- | --- | --- | --- |
| AIR | 350–750 | 0.08 | 0.98 | 0.98 |
| SY60WF IOL | 453.41–749.75 | 0.13 | 0.45 | 0.45 |
| SING IMT<sup>TM</sup> | 397.08 – 749.75 | 0.12 | 0.93 | 0.91 |

### Geometrical aberrations

Figure 4A shows the wavefront of the SING IMT™ reconstructed as the linear combination of Zernike polynomials. A clear curvature of the wavefront is evident with a concavity indicating that the SING IMT™ behaves as a diverging lens with a focal length of about -100 mm. The expansion of the wavefront into a Zernike basis allowed us to quantify the extent of the first 14 aberrations. As shown in Figure 4B and Table 2, the SING IMT™ has negligible coma, trefoil, astigmatism, and spherical aberrations, which clearly proves its suitability as optical element. In the same *in-vitro* conditions, the SY60WF monofocal IOL behaves as a converging lens with a focal length of ∼ 45 mm, which is in agreement with the characteristics reported by the manufacturer and the clinical purpose of this intraocular lens, namely, replacing the crystalline in cataract surgery. Yet, the normalized Zernike coefficients are slightly larger with respect to those obtained with the SING IMT™ device (Figure 4C-D), which might be ascribed to possible mechanical stress that the flexible haptics transfer to the lens in the *in-vitro* conditions of our experiments. Note that the SING IMT™ lens does not suffer from deformation-induced geometrical aberrations because of its higher mechanical rigidity.

**Table 2:** Absolute values of Zernike coefficients of the SING IMT™

| Aberrations | SING IMT™ [$\mu\text{m}$] |
| --- | --- |
| Tip Y | -0.057 |
| Tilt X | -2.652 |
| Astigmatism $\pm 45^\circ$ | -0.034 |
| Defocus | -7.247 |
| Astigmatism $0^\circ / 90^\circ$ | - 0.028 |
| Trefoil Y | 0.005 |
| Coma X | -0.007 |
| Coma Y | -0.004 |
| Trefoil x | -0.009 |
| Tetrafoil Y | 0.025 |
| Sec. Astig. Y | 0.003 |
| Spher. Aberr. 3 | -0.011 |
| Sec. Astig. X | -0.016 |
| Tetrafoil X | 0.017 |

**Figure 4:**
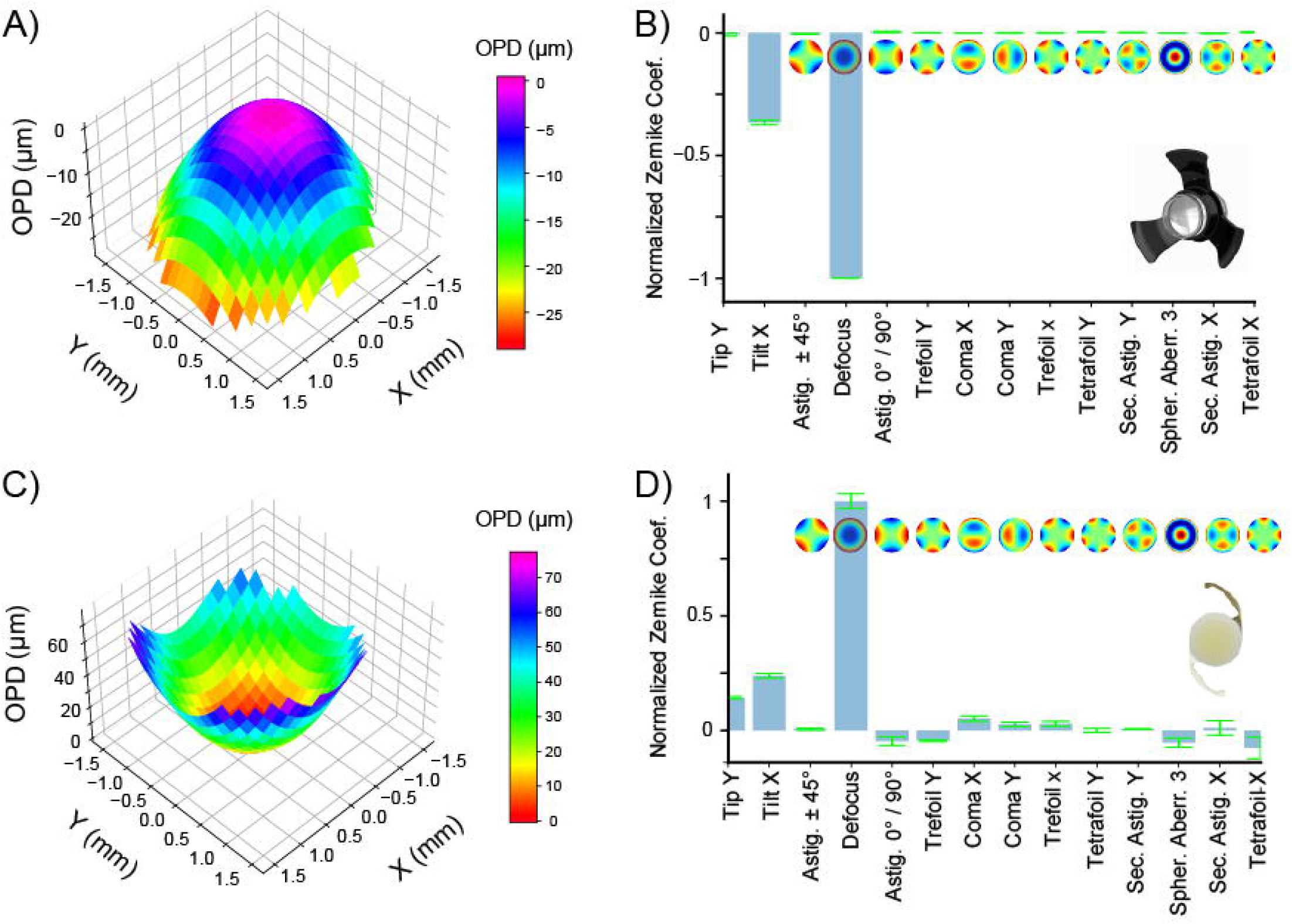
Wavefront sensing and geometrical aberrations characterization. A) Reconstructed wavefront and B) Zernike coefficients of SING IMT™ device. C) Reconstructed wavefront and D) Zernike coefficients of SY60WF monofocal IOL. For clarity of comparison, the Zernike coefficients of each lens are normalized to the absolute value of the defocus term. The optical path difference (OPD) is defined as the difference between the aberrated and the ideal unaberrated wavefronts.

## Discussion

Intraocular vision-improving devices, such as implantable lenses, have been recently used to recover the vision of patients affected by various forms of AMD.^12^ However, AMD-affected patients usually have poor functional results after cataract surgery with standard monofocal IOL implants.^13^ To overcome this problem, novel and more advanced intraocular devices, such as miniaturized telescopes, have been developed.^14-16^ For instance, the IMT™ (Vision Care, Israel), which is the precursor of the SING IMT™ and the first evaluated in a large and prospective 2-year multicenter pivotal study with more than 200 patients, was proved effective in improving the visual acuity of patients with moderate to profound visual impairment caused by bilateral end-stage AMD.^17^ Following these promising results, the IMT™ was further developed leading to the creation of the SING IMT™, which is currently approved by the US Food and Drug Administration (FDA), Canadian, and European authorities as a monocular implant for patients aged at least 55 years with stable and severe vision impairment (BCDVA 20/80 to 20/800) caused by bilateral central scotomas due to end-stage AMD.^18^

The operational principle of the SING IMT™ is quite simple. Briefly, the SING IMT™ is an ultra-precision wide-angle micro-optic whose magnification compensates for missing scarred macular receptors, thus attenuating the visual impact of scotoma. Though proven functional, the quality of the vision recovered with SING IMT™ might be sensitive to its optical features, as a compound optical system. To erase possible doubts on this point, here we assessed the optical performance of the SING IMT™ device under *in-vitro* experimental conditions. In particular, we studied the spectral transmission of the SING IMT™ in the visible spectrum and measured its geometrical aberrations. A commercial intraocular lens, namely the SY60WF monofocal IOL, Alcon (branded as Clareon IOL), was used as a benchmark for this study.

The SING IMT™ lens exhibited a quite flat transmission spectrum over the visible interval (i.e., 400-750 nm). Furthermore, the SING IMT™ device and SY60WF monofocal IOL feature high UV-rejection and blue-rejection capabilities, respectively. These results agree with the optical properties of the structural materials of the two optical elements, namely polymers for the SY60WF monofocal IOL^19-21^ and fused silica for the SING IMT™.^22-24^ Moreover, the flatness and large bandwidth of the SING IMT™ suggested the absence of resonance between the lenses of the compound system due to spurious or undesired reflections.

The wavefront’s concavity indicated that the SING IMT™ behaves as a diverging lens with a focal length (absolute value) ∼2.4-folds higher than that of the crystalline lens^25^, which is in agreement with the magnification expected once implanted into the eye, which is 2.7 ± 10% as by design. Moreover, wavefront sensing and Zernike analysis revealed that both devices are suitable as intraocular imaging elements. In particular, the SING IMT™ showed excellent optical quality with almost zero coma, astigmatism, trefoil, and spherical aberrations. The only terms that contribute to the wavefront shape are tilt-X ad defocus. However, it should be noted that defocus and tilt are not real aberrations, since they only shift the focal plane. In other words, a perfect wavefront altered by tilt or defocus still result in an aberration-free image. Notably, this result can be ascribed to the design of the SING IMT™, where the use of flexible polymers is limited to the haptics. The latter are indeed the only elements that could introduce tilting of the wavefront because the optical element is a rigid compound system made of silica microlenses. As such, this design prevents mechanical deformation of the optics^26^ and, thus, formation of geometrical aberration, which is a key aspect for high-quality imaging.

Importantly, although the SING IMT™ has excellent optical quality it is still crucial to follow strict patient eligibility criteria to obtain a functional result.^18^ Precisely, patients should be advised of the complete loss of stereopsis that results from the lens implant, which is typically done only in one eye, so that the other eye can be still used to see in the surrounding space while the implanted eye is fitted for near vision. For this reason, after surgery multiple rehabilitation sessions are mandatory to avoid diplopia and allow proper use of the implanted optical device, regardless of its high optical quality.

## Conclusions

In this work we assessed the optical quality of the intraocular miniaturized compound lens system, namely the SING IMT™ device (Samsara Vision), through an *in-vitro* characterization of its geometrical aberrations and optical transmittance in the UV-Vis. We found that the SING IMT™ effectively operates as a diverging lens with homogenous high-optical transmittance in the visible along with UV-rejection. Precisely the SING IMT™ is capable of magnifying the retinal images without compromising optical quality making it a valuable choice in advanced-stage AMD patients undergoing cataract surgery.^13^ Moreover, the SING IMT™ exhibited a regular wavefront as confirmed by Zernike’s analysis, which resulted in negligible geometrical aberrations. The feasibility of the SING IMT™ as an intraocular implant is further evident if compared with intraocular technologies entirely based on flexible polymers, at least under *in-vitro* experimental conditions.

## Acknowledgements

This work is partially supported by Samsara Vision, which also provided the intraocular devices for the experimental measurements.

## Conflicts of interest

Dr. Faustino Vidal-Aroca reports also being an employee of Samsara Vision during this study. The other authors declare no conflict of interest. The funders had no role in the design of the study; in the collection, analyses, or interpretation of data; in the writing of the manuscript; or in the decision to publish the results.

