## Supplemental Materials for "Optical-quality assessment of a miniaturized intraocular telescope"

Running head: Miniaturized intraocular telescope

Authors: Irene Nepita PhD[a,b*], Raffaele Raimondi MD[c], Simonluca Piazza PhD[a,b], Alberto Diaspro PhD [a,b], Faustino Vidal-Aroca PhD[d], Salvatore Surdo PhD[b,e*], Mario R. Romano MD PhD[c,f]

Affiliations:

[a] Nanoscopy&NIC@IIT, Istituto Italiano di Tecnologia, Genoa, Italy
[b] Genoa Instruments s.r.l., Genoa, Italy
[c] Department of Biomedical Sciences, Humanitas University, Milano, Italy
[d] Medevise Consulting, 67000 Strasbourg, France
[e] Dipartimento di Ingegneria dell'Informazione, Università di Pisa, Pisa, Italy
[f] Eye Center, Humanitas Gavazzeni-Castelli, Bergamo, Italy

Corresponding author:
Dr. Salvatore Surdo
Dipartimento di Ingegneria dell'Informazione
Università di Pisa
Pisa, Italy

Phone number: +39 050 2217 504

**Keyword list:** End-stage age-related macular degeneration; visual impairment; visual prosthesis; implantable ophthalmic device; intraocular lens; optical performance, geometrical aberrations; small-incision new generation implantable miniature telescope SING IMT™.

### Sample holders engineering

In order to guarantee high precision in the alignment of the optical lens under test to the probing light, we designed and prepared custom mounts and cage systems that take into account the overall dimensions and geometries of the intraocular devices. For the SING IMT™ device, an adjustable circular iris diaphragm (SM05D5D, Thorlabs) was mounted into a 30 mm cage plate with removable sample holder (CFH1R, Thorlabs) (Figure S1). The implantable device was positioned in the iris aperture and the three haptic wings were blocked with a rubber O-ring, avoiding additional forces and, thus, limiting possible off-axis misalignments.

We could not use this sample holder for the monofocal intraocular lens (SY60WF, Alcon) because of its elasticity, which might cause undesired deformations under the action of uneven mechanical pressure. For this reason, a sample holder compatible with standard mounting systems of 1-inch optics (Figure S2) was designed and 3D printed.

**FIGURE S1**

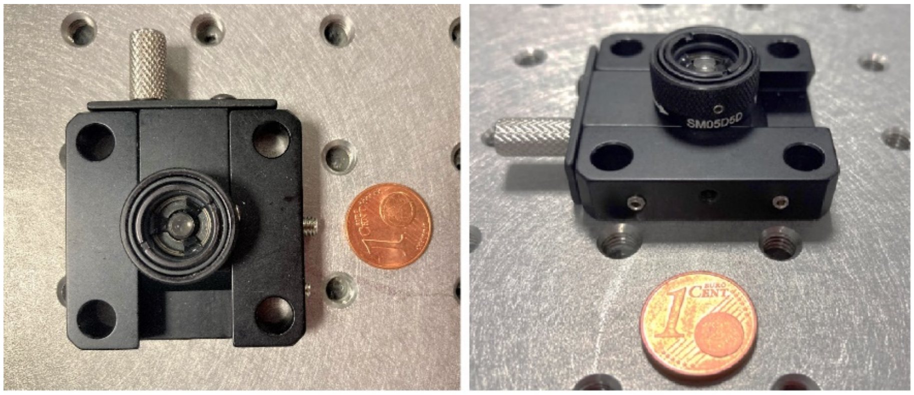


*Figure S1. SING IMT™ device holder. Pictures showing the sample holder mounting the SING IMT™ and used to precisely insert the optical element under test into the measurement setup.*

**FIGURE S2**

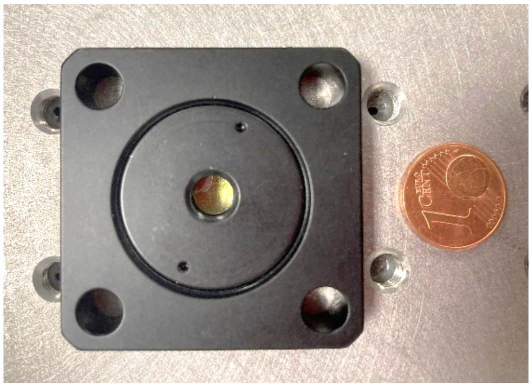

*Figure S2: monofocal intraocular lens (SY60WF, Alcon) device holder. Picture of the 3D printed sample holder for the IOL lens.*

**FIGURE S3**

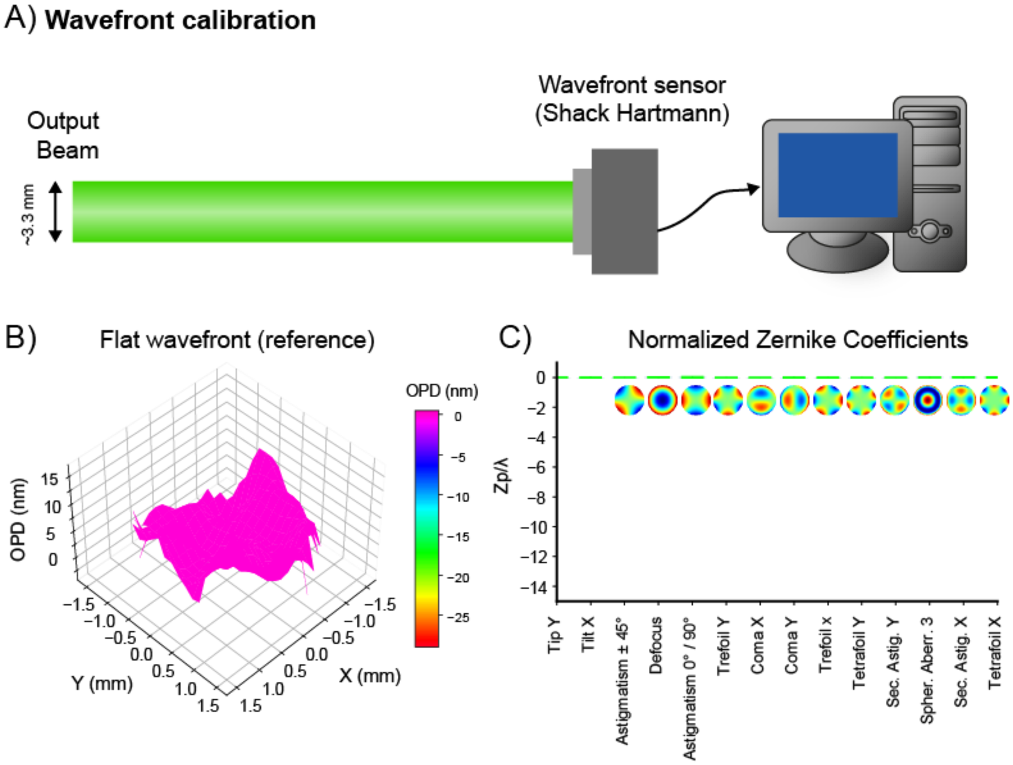


*Figure S3: Calibration of the wavefront sensing setup. A) Schematic description of the optical setup used to calibrate the wavefront sensor. B) Measured reference wavefront and corresponding geometrical aberrations C). The optical path difference (OPD) is defined as the difference between the aberrated and the ideal unaberrated wavefronts.*
